# Synthetic negative feedback circuits using engineered small RNAs

**DOI:** 10.1101/184473

**Authors:** Ciarán L. Kelly, Andreas W. K. Harris, Harrison Steel, Edward J. Hancock, John T. Heap, Antonis Papachristodoulou

## Abstract

Negative feedback is known to endow biological and man-made systems with robust performance in the face of uncertainties and disturbances. To date, synthetic biological feedback circuits have relied upon protein-based, transcriptional regulation to control circuit output. Small RNAs (sRNAs) are non-coding RNA molecules which can inhibit translation of target messenger RNAs (mRNAs). In this paper, we designed, modelled and built two synthetic negative feedback circuits that use rationally-designed sRNAs for the first time. The first circuit builds upon the well characterised *tet*-based autorepressor, incorporating an externally-inducible sRNA to tune the effective feedback strength. This allows more precise fine-tuning of the circuit output in contrast to the sigmoidal input-output response of the autorepressor alone. In the second circuit, the output is a transcription factor that induces expression of an sRNA which negatively regulates the translation of the mRNA encoding this output, creating direct, closed-loop, negative feedback. Analysis of the noise profiles of both circuits showed that the use of sRNAs did not result in large increases in noise. Stochastic and deterministic modelling of both circuits agreed well with experimental data. Finally, simulations using fitted parameters allowed dynamic attributes of each circuit such as response time and disturbance rejection to be investigated.

## INTRODUCTION

Natural biological systems function through the regulation of a large number of genes and their products, which interact with each other dynamically and precisely in response to a wide variety of signals. The ability to engineer biological systems to perform such complex tasks is a key goal of synthetic biology (1), with limited success to date (2, 3). A number of engineering approaches have been adopted in the design and implementation of synthetic biological circuits (4–11) yet many challenges remain. Chief amongst them is a lack of performance reliability in novel, untested contexts and when dynamic disturbances to the circuits are applied (12).

Employing negative feedback control is known to endow systems with robustness and improved performance. This is particularly valuable when prediction of a circuit component’s behaviour in a particular context is impossible, and when disturbances to the system are applied that alter the performance of these components (2, 13–17). Increased robustness of output also allows the reliable connection of two or more circuits where the output forms the input of the second circuit (12, 16). The basic architecture of a feedback circuit is as follows: an input is applied to a system, in order to regulate its output to a desired value; this output is measured and compared to the desired value; if the measured output is different to the desired one, an error signal results; this error signal is fed back to the input of the system, thereby pushing the output towards the desired value. The object responsible for responding to the error signal is called the “controller” and the point where the external and “fed-back” inputs are combined is called a “summing junction”. Negative feedback is predominantly used to reduce variation in the output of a system. The addition of well-characterised, insulated feedback control architectures to biological systems can overcome dynamic fluctuations in the copy number and activity of biological parts such as genes, mRNAs and proteins, and ensuring a consistent output. The output species could be a protein, a nucleic acid or even a small molecule. Indeed, negative feedback motifs are so resilient that function can be maintained despite mutations in circuit components, allowing greater evolutionary flexibility (18). One potential side-effect associated with the use of feedback systems is the introduction of additional noise into the system, resulting in increased variability of the output. This additional noise can be introduced during sensing operations and at summing junctions.

Unsurprisingly given the many advantages its use confers, feedback is ubiquitous in natural biological systems (19–27). *In vivo* feedback has been employed in many synthetic gene circuits, including circuits such as the toggle switch (28, 29), the repressilator (30), sustained and tunable oscillators (31) and a concentration tracker (32). *In silico* feedback has also been successfully employed to control gene expression (33), with improved understanding leading to more successful synthetic feedback circuits (34). Most synthetic biological feedback circuits constructed to date have used a protein-based controller and can be divided into three categories: circuits that reduce the concentration of a protein through sequestration or increased degradation, where transcription is the process under control and a protein-protein summing junction is used (32, 35); transcriptional feedback circuits where transcription is controlled and a protein-DNA summing junction is used (28, 30, 31, 36, 37); and finally translational feedback circuits where translation is controlled and a protein-mRNA summing junction is used (38). One widespread and well-studied transcriptional negative feedback architecture is the negative autoregulator or autorepressor, found both in natural (22, 39–47) and synthetic circuits (22, 39, 40, 42, 48). An autorepressor consists of a transcription factor (TF) which binds to the promoter responsible for expression of the TF gene, repressing transcription and reducing the concentration of the TF in the cell. The use of autorepressor architectures in synthetic biology is however hindered by the steep, sigmoidally-shaped input-output curve, allowing only coarse tuning of the regulated output with an external input (49). One way to overcome this problem could be to adjust the feedback strength. This can be achieved through: scaling the feedback signal by using weaker ribosome-binding sites (RBSs), engineering of the transcription factor to alter its inducer and promoter interactions, mutagenesis of the promoter to change transcription-initiation rates, or increasing the low degradation rate of the transcription factor (50, 51). All four of these solutions are hard-wired into the circuit and not easily manipulated during cell growth. Additionally, higher turnover of protein consumes cellular resources such as ribosomes, rRNAs and amino acids, increasing the burden of the circuit on the cell.

Post-transcriptional or translational feedback through the use of RNA controllers and an RNA-RNA summing junction is an appealing alternative to transcriptional feedback via protein-based controllers. Translational regulation reduces the effect of leaky transcription resulting in more tightly-controlled feedback than transcriptional regulation where mRNAs continue to produce protein until degraded (38). Theoretical work suggests that translational feedback mitigates fluctuations in translational resources that could destroy circuit performance (52). Increased understanding of natural regulatory RNAs (53, 54), coupled with the predictability of the folding and interactions of RNA structures, has led to a resurgence of interest in RNAs as a means of regulation (55, 56). Functional small RNAs are faster for a host cell to produce and require less energy than protein production (45), constituting a lower burden to the host, a key consideration for synthetic biological circuits (57). They also have the potential to propagate signals rapidly (49, 58, 59), as their dynamics are tied to the naturally high degradation rate of RNA. Hfq (host factor for RNA phage Q beta replication)-associated sRNAs are found in many prokaryotes (60, 61), are involved in a wide range of host cellular functions (53, 54, 62–64) and have been studied extensively at the biochemical level (54, 65–68). The regulation of translation by sRNAs can occur through one of four mechanisms: inhibition of translation; activation of translation; stabilisation of the duplexed mRNA; or stimulation of degradation of the target mRNA (53, 54, 62, 69). The inhibition of mRNA translation by sRNAs has great potential for use in synthetic negative feedback circuits and has been explored theoretically (70–72) and using the analogous mechanism of RNAi in mammalian cells (73). The non-feedback use of synthetic sRNAs against a small number of targets has been reported (74–76) and design principles described (77). The mechanism of inhibition has been modelled (59, 78, 79) and compared to experimental data (49, 80–82). As synthetic sRNAs operate primarily through complementarity, they can be designed to target any sequence, allowing the construction of a greater diversity of feedback circuits than currently possible using transcription factors alone. Additionally the input-output response curves of sRNAs have been shown to be almost linear, unlike the steep, non-linear response curves of repressing transcription factors (49), which could be useful to tune feedback circuits. The use of sRNAs controllers would be particularly useful when large circuits involving many proteins are to be connected together, removing the burden of another dynamic and highly produced protein (16). One potential drawback of sRNA-based feedback circuits is that due to their high degradation rates, sRNAs are often noisier than their protein-based counterparts (59). Translational feedback however, is predicted to lower intrinsic noise (83, 84).

In this work we demonstrate two novel uses of Hfq-associated sRNAs in synthetic negative feedback circuits. In the first circuit we take the well-studied TetR-based autorepressor and through simulations and experimental data, we demonstrate that it is possible to tune the effective feedback strength of an autorepressor via the inducible expression of a translation-inhibiting sRNA (Figure 1A). The ability to externally and temporally fine-tune the effective feedback strength allows easier and more precise access to the full range of steady-state outputs than with the TetR-based autorepressor alone. In the second circuit we construct a closed-loop negative feedback circuit using sRNA instead of a repressor protein to control the translation of an activating-transcription factor which is responsible for expression of the sRNA (Figure 1B). The performance of this circuit is modelled and tested experimentally and the ability to set the output steady state and tune feedback strength using two external inputs is demonstrated. The intrinsic and extrinsic noise of both circuits is also simulated and measured experimentally with no increase in noise with increasing sRNA concentrations observed. Finally using experimentally-derived parameters, predictions were performed of the dynamic responses and disturbance rejection properties of each circuit, highlighting further benefits of sRNA-based negative feedback circuits.

**Figure 1.**
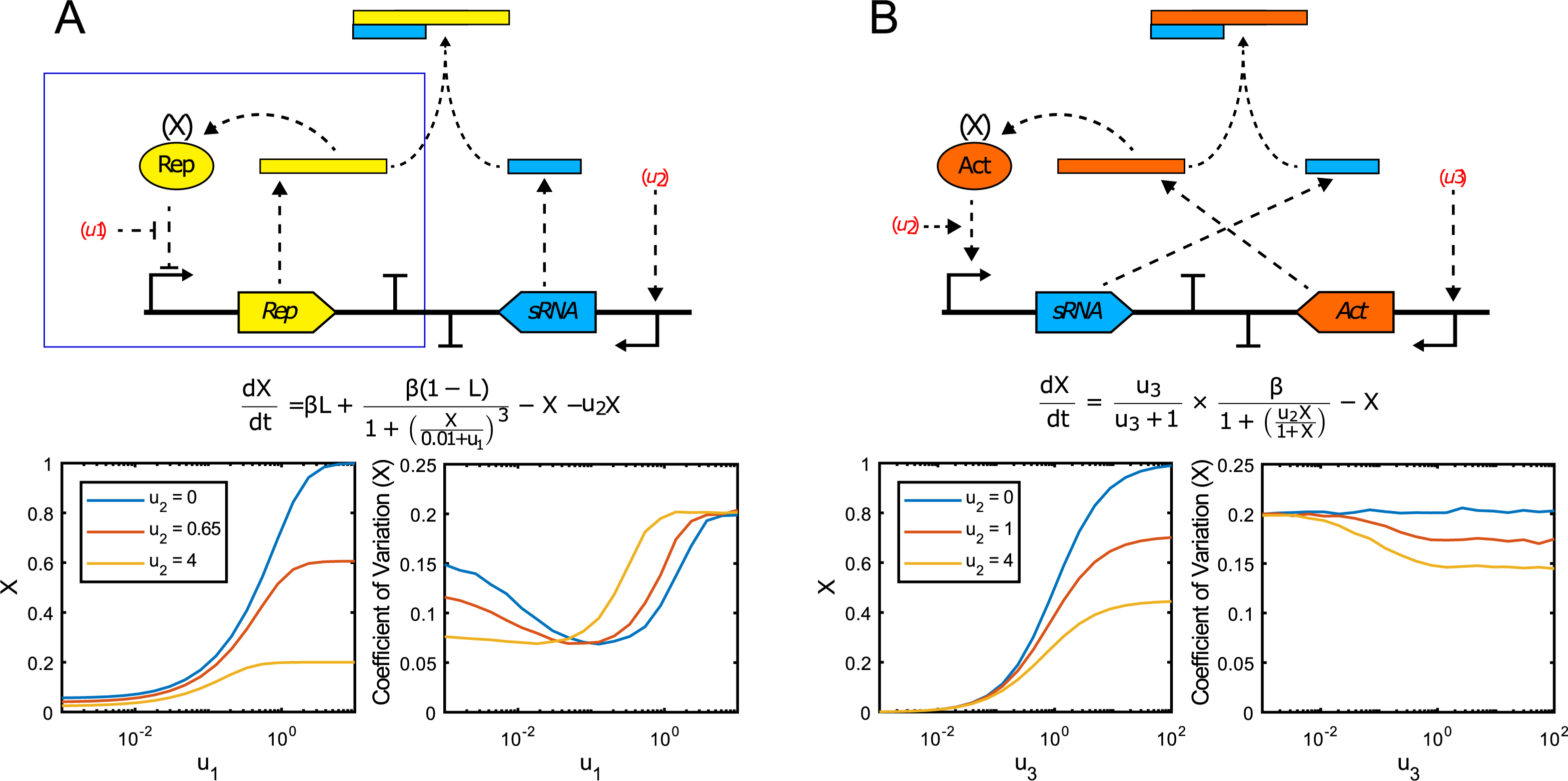
Schematic diagrams of the circuit architectures investigated in this study and modelling predictions of each circuit’s performance. A. An sRNA-tuned autorepressor circuit consisting of: a gene encoding a repressor protein (Rep); a promoter upstream of the repressor-encoding gene *(Rep)*, which the repressor protein can bind to and inhibit transcription from; an sRNA, whose expression can be induced through an external input (*u*_2_) and which reduces the translation of the autorepressor mRNA, thus reducing the effective feedback strength of the autorepressor loop (blue box). Simulations predict that the addition of the sRNA would increase the fidelity of steady-state output (X) tuning through the use of two external inputs (*u*_2_ and *u*_1_; red and yellow lines) instead of one (*u* only; blue line). Additionally the use of an sRNA should not result in large increases in output noise. B. A closed-loop sRNA feedback circuit consisting of a translation-inhibiting sRNA, whose expression can be induced through an external input (*u*_2_), in closed-loop feedback with an mRNA encoding the transcriptional activator (Act) of the inducible promoter upstream of the sRNA-encoding gene. The steady-state output can primarily be set using a different external input (*u*_3_) which binds to a constitutively-expressed transcriptional activator (not shown) inducing transcription of the *Act* gene. Simulations predict that the closed sRNA-act mRNA feedback loop would reduce the range of the steady-state output (area between blue and yellow lines). Additionally the use of an sRNA is predicted to reduce output noise.

## MATERIALS AND METHODS

### Mathematical modelling

The synthetic negative feedback circuits outlined in this paper were simulated using mathematical modelling (Supplementary Information Section 1). Regulation at both the trancriptional (repressor-promoter) and translational (sRNA-mRNA) levels were first modelled using simplified descriptions of the biochemical processes involved with nominal parameter values, allowing *a priori* qualitative prediction of the various circuits’ behaviour (Figure 1). The autorepressor was then simulated using a two-state model (SI 1.3) similar to previously described models of the TetR autorepressor (85). Experimental data was used to identify the unknown parameters in the model using least-squares minimisation, and the model fit compared to the experimental results. The autorepressor model was extended via inclusion of two extra states to account for sRNA inhibition of translation, and experimental data was again used to identify the unknown parameters. The fitted model was then used to perform dynamic simulations of the sRNA-tuned autorepressor circuit’s response to an input disturbance. Finally, the closed-loop sRNA negative feedback circuit was simulated using a four state model (SI 1.5) and fit to the experimental results and the model was again used to simulate the circuit’s dynamic response to changing inputs. The fit models are then used to estimate the influence of both intrinsic noise (via stochastic simulations) and extrinsic noise (via parameter variation modelling) on the variability of each circuit’s output. These sources are then combined and compared to experimental measurements of Coefficient of Variation (CoV).

### Bacterial Strains and Growth Conditions

*E. coli* strain DH5α was used for all plasmid construction and propagation. Wild-type *E. coli* strain MG1655 was used for initial experiments with pCK200 - pCK209 plasmids. Subsequently the Keio collection *rhaS* and *rhaBAD* mutant JW3876 was used for all further time course assays. Cells were cultured in either LB, M9 minimal media supplemented with 0.5 % v/v glycerol, or EZ rich defined media (Teknova Inc) supplemented with 0.5 % v/v glycerol and incubated at 37 °C with vigorous shaking. Ampicillin and chloramphenicol were used at final concentrations of 125 μg/mL and 25 μg/mL respectively.

### Plasmid Construction

A table of plasmids and oligonucleotides used in this study is provided in Table S3. Sequences of all plasmids have been submitted to Genbank. All plasmid construction was carried out using standard molecular cloning methods. Full details are provided in Supplementary Information Section 2. All synthetic DNA fragments (gblocks) were synthesised by Integrated DNA Technologies Inc.

### Assays

*E. coli* MG1655 or JW3876 (*ΔrhaS* Keio collection mutant, *ΔrhaBAD* ensuring no metabolism of L-rhamnose) cells were transformed with appropriate plasmid(s) and isolated transformants used to inoculate 5 ml of EZ rich defined media (rich media hereafter) or M9 minimal media (minimal media hereafter) supplemented with 0.5 % (v/v) glycerol, as appropriate, and ampicillin or chloramphenicol, which was cultured with shaking at 37 °C for 16 h. For flow cytometry experiments, these cultures were then subcultured into 300 μL of fresh media in deep 96 well multiplates, to an optical density (OD; A600 nm) of 0.05 and the plate incubated at 37 °C with rapid shaking (700 rpm). For circuits using the *tet* promoter, the tetracycline analogue, anhydrotetracycline (aTc) was added to the assay cultures at the concentrations specified. For circuits using the *rhaBAD* promoter, the inducer L-rhamnose was added to the assay cultures at the concentrations specified. For circuits using the *Pm* promoter, the inducer *m*-toluic acid was added to assay cultures at the concentrations specified. Samples were taken at specified time points for optical density and GFP fluorescence measurements. A FLUOstar Omega platereader (BMG Labtech) was used for all absorbance measurements of optical density (OD) at 600 nm and GFP fluorescence measured using an Attune NXT flow cytometer (ThermoFisher Scientific). Cells were gated using forward and side scatter, and GFP fluorescence (excitation and emission wavelengths: 488 and 525 nm [with 20 nm bandwidth] respectively) was measured. Histograms of fluorescence intensity were plotted, and geometric mean statistics extracted. Coefficients of variation were extracted from the gated population as follows. FCS files were created from the gated cells from flow cytometry and raw reads extracted using the FCS Extract Utility created by Earl F. Glynn at the Stowers Institute for Medical Research. The resulting log-normally distributed data was analysed using equation 10 (S.I. 1.4) to calculate the CV. For platereader experiments, overnight cultures were subcultured into 150 μL of fresh media in 96-well microplates, to an optical density (OD; A600 nm) of 0.05 and the plate incubated at 37 °C with rapid shaking in a FLUOstar Omega platereader (BMG Labtech). Optical density and GFP fluorescence (excitation and emission wavelengths: 485 and 530 nm [with 20 nm bandwidth] respectively) were measured every 15 minutes, with fluorescence normalised by optical density and plotted over time.

## RESULTS

An externally-regulated sRNA to tune the effective feedback strength of an autorepressor The use of a translation-inhibiting sRNA to reduce the effective feedback strength of the autorepressor in order to tune its output was explored. This solution is appealing as it can be bolted onto any existing transcription-based feedback loop, avoiding modification of the DNA encoding the core feedback circuit (as would be required to modify the scaling of the feedback signal using RBSs of different strengths). As a starting point, a plasmid encoding an autorepressor based on the well-studied tet-repressor mechanism (22, 39–42) was constructed. The plasmid consists of the *tet* promoter (P_iei_) which controls the transcription of a gene encoding the TetR repressor, which can bind to P_*tet*_ reducing *tetR* transcription. The tetracycline analogue, anhydrotetracycline (aTc) is an inducer of the P_*tet*_-TetR system, preventing TetR binding to the promoter, thereby increasing expression from P_*tet*_. This provides one external input to the system with which to change the output steady state. In our system the *tetR* gene is fused to the gene encoding superfolder GFP (sfGFP) allowing easy quantification of the TetR protein concentration. A strong, synthetic RBS with a translation initiation rate (T.I.R.) of 50,000 was designed and inserted upstream of *tetR*, and a plasmid backbone with a copy number of 15-20 chosen. The performance of the resulting plasmid, pCK200 (Figure S2A), was tested experimentally.

The wild-type *E. coli* strain MG1655 containing pCK200 was grown in both minimal and rich media and its fluorescence monitored over time. Its dynamic response in minimal media (Figure S2B) was found to be in good agreement with previously described autorepressor circuits (40). The sigmoidal response of the steady-state output (measured in cells grown in rich media at exponential phase of growth; Figure S14) to increasing concentrations of the input (*u*_1_,), anhydrotetracycline (aTc) is clearly demonstrated (Figure S2C), highlighting the difficulty in tuning the output of circuit. Lastly, unknown parameters in the system were identified from the experimental data. With these parameters, the model was found to accurately capture the system’s behaviour over the range of inducer concentrations used, and correctly approximate the low- and high-inducer saturation levels anticipated from this promoter (Figure S3).

Having validated the autorepressor module, we next set about designing a translation-inhibiting small RNA (sRNA) that targets the tetR-sfGFP mRNA of the autorepressor feedback loop. The Hfq-binding scaffold of the *E. coli* micC sRNA and the strong Rho-independent T1/TE terminator were chosen, based on previously-described design principles (74, 75, 77). In order to maximize translation inhibition of the target mRNA, two different targeting sequences, each 24 nucleotides in length, were designed and tested. The first sequence was designed to bind the first 24 nucleotides of the tetR-sfGFP open reading frame (named “Start” hereafter) and the second designed to bind 24 nucleotides including all of the probable Shine Dalgarno sequence upstream of the tetR-sfGFP open reading frame (named “SD” hereafter). Both sequences were computationally analysed for binding to the tetR-sfGFP mRNA and for off-target binding to the wild-type *E. coli* (MG1655) ribonucleome using IntaRNA (86–88). Binding of the SD sRNA to the tetR-sfGFP mRNA (−37.35 kcal/mol) was predicted to be more energetically favourable than that of the Start sRNA (−31.09 kcal/mol), implying stronger binding. Binding to off-target mRNAs was substantially less favourable than either of the target sequences (Table S2) and therefore considered unimportant. Each of these sRNA constructs were inserted into the backbone of a medium copy (15-20 copies per cell) *E. coli* expression plasmid and a strong constitutive promoter (P_proD_) inserted upstream of either, resulting in plasmids pCK209 (SD) and pCK220 (Start). Next the ability of these sRNA plasmids to inhibit translation of the tetR-sfGFP mRNA was tested. *E. coli* strain MG1655 containing pCK210 (identical to pCK200 but with *rhaS* inserted downstream of *ampR*; required for inducible expression of sRNA in following section) and either pCK209, pCK220 or pAH23 (empty vector control; no sRNA) were grown in rich media supplemented with different concentrations of aTc until late exponential phase (5 h; Figure S14) and fluorescence measured by flow cytometry. Both SD and Start sRNAs were extremely effective in shutting down tetR-sfGFP mRNA translation from the autorepressor with no aTc added (highly statistically-significant difference; *P* < 0.0001) (Figure 2B) and showed similar translation-inhibition of tetR-sfGFP mRNA when aTc was added (Figure 2A). Encouragingly the use of an sRNA to reduce the output from the autorepressor circuit did not result in large changes in the noise profile of the circuit, with the response of the coefficient of variation (a relative measure of the variability around the mean) to increasing concentrations of aTc similar in all three sample groups (Figure 2C). The SD-targeting sRNA was taken forward for all subsequent experiments in this circuit, based on the lower predicted binding energetics.

**Figure 2.**
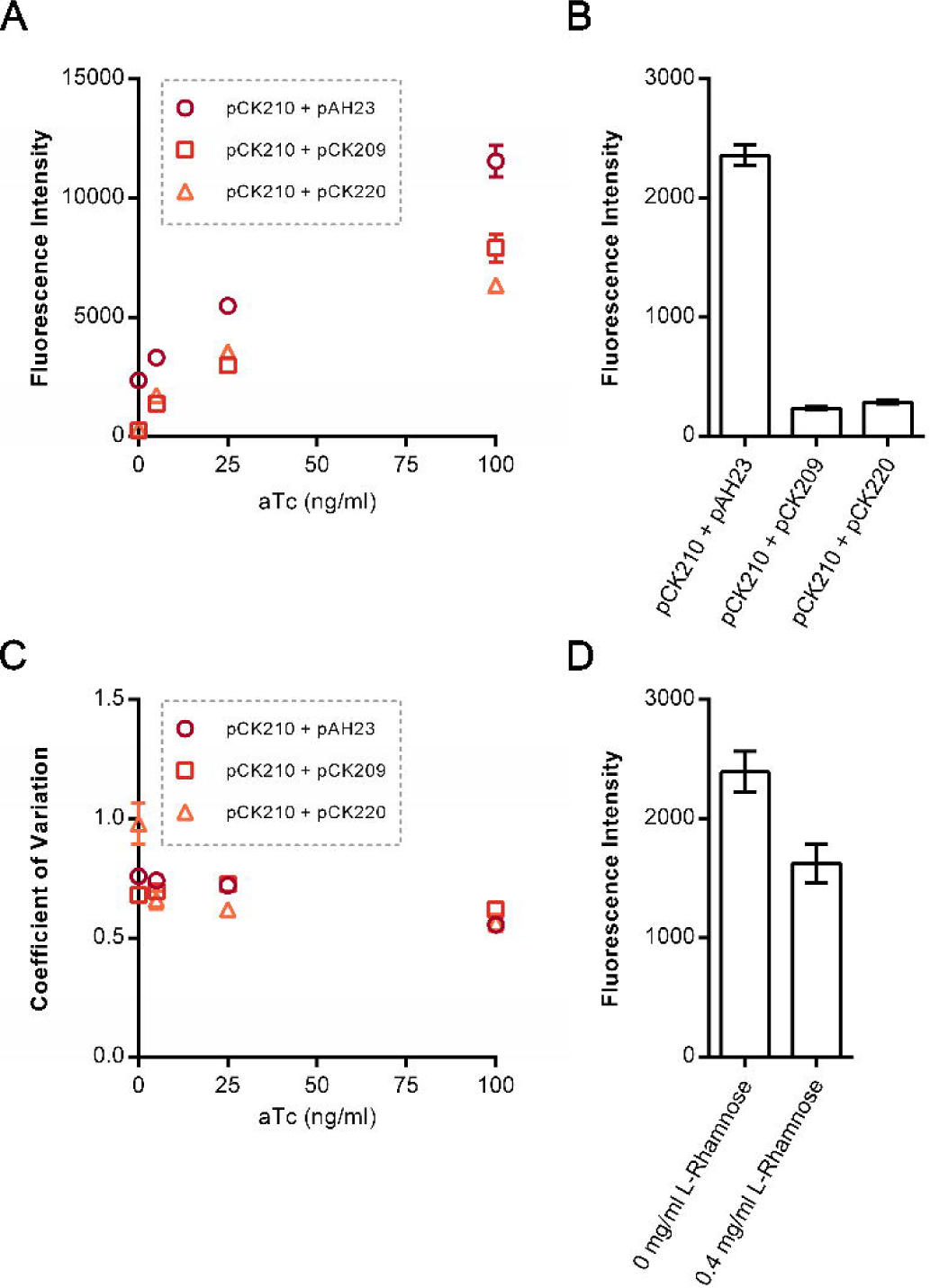
Testing of two synthetic sRNAs targeting the tetR-sfGFP mRNA, and investigating the inducibility of an sRNA. A. Plasmids encoding sRNAs targeting the Shine Dalgarno (pCK209) or start codon (pCK220) of the tetR-sfGFP mRNA were designed and constructed and the ability of each sRNA to inhibit translation assessed. *E. coli* strain JW3876 was transformed with pCK210 (encoding the autorepressor) and one of pCK209, pCK220 or pAH23 (negative control, no sRNA), cultured at 37 °C in EZ rich defined medium supplemented with glycerol and increasing concentrations of aTc, and GFP fluorescence measured at late exponential phase (5 h) by flow cytometry. B. Comparison of the 0 ng/ml aTc samples from A. C. Experimentally-obtained coefficients of variation around the TetR-sfGFP mean from A. D. To test the inducible expression of sRNA,the proD promoter of pCK209 was replaced with the *E. coli rhaBAD* promoter resulting in plasmid pAH17. *E. coli* strain JW3876 was transformed with pCK210 and pAH17, cultured at 37 °C in EZ rich defined medium supplemented with glycerol and 0 or 0.4 mg/ml L-rhamnose, and GFP fluorescence measured at late exponential phase (5 h) by flow cytometry. Fluorescence intensity represents the geometric mean of fluorescence. Error bars shown represent the standard deviation of three independent biological replicates.

We next explored the tunability of the sRNA inhibition system, by replacing the static constitutive promoter with an inducible promoter. This introduces a second external input to the autorepressor feedback loop, allowing the ratio of sRNA to mRNA to be varied precisely. In contrast to other approaches to alter the steady state output, such as changing the RBS of the tetR-sfGFP mRNA to directly scale the feedback signal, the use of an sRNA dial does not require changing the DNA encoding the central autorepressor circuit. The well-characterised *E. coli rhaBAD* promoter (P_*rhaBAD*_) (89, 90) was inserted in place of the constitutive promoter in front of the sRNA construct, resulting in plasmid pAH17. The Keio collection *E. coli* mutant, JW3876 (91) was used to test the performance of this inducible sRNA plasmid. This strain lacks *rhaS*, the transcriptional activator of *P*_*rhaBAD*_, and the entire *rhaBAD* operon, encoding the enzymes for L-rhamnose consumption and thus should ensure maximal insulation from crosstalk with host metabolism. JW3876 cells containing both pCK210 and pAH17 were cultured in rich media with/without 0.4 mg/ml L-rhamnose to mid-exponential phase (4 h; Figure S14) and fluorescence analysed by flow cytometry (Figure 2D). As expected the level of fluorescence was significantly lower with the addition of L-rhamnose (statistically-significant difference; *P* < 0.01). It was surprising that even with such a high concentration of inducer the level of translation inhibition was less than with either of the two constitutive promoter sRNA plasmids. Further inspection of *rhaBAD* promoter revealed the inclusion of five additional bases after the transcriptional start site (TSS +1), an artefact of misannotation of the promoter that could affect sRNA-mRNA interaction. Additionally when spread across two plasmids, the autorepressor and sRNA constructs would be at different copy numbers.

In response to both of these observations, the minimal *rhaBAD* promoter (lacking the unnecessary bases downstream of the TSS) and the SD sRNA-encoding fragment was inserted into pCK210, resulting in pCK221 (Figure 3A). JW3876 cells containing pCK221 were grown in rich media supplemented with increasing concentrations of L-rhamnose and no aTc and the fluorescence of cells measured at late exponential phase (5 h; Figure S14). Mean fluorescence is reduced with increasing concentrations of L-rhamnose, with saturation observed at 0.001 mg/ml L-rhamnose (Figure 3B). This concentration corresponds with saturating levels of the RhaS-P_*rhaBAD*_ induction system and agrees with data from strains incapable of L-rhamnose metabolism or when a non-metabolisable inducer of L-rhamnose is used (89, 92). This result confirmed the hypothesis that varying the concentration of sRNA in the cell would allow the output of the autorepressor circuit to be varied independently of aTc. We next sought to test the hypothesis that the use of the sRNA dial and the second input (*u*_2_; L-rhamnose), and the first input (*u*_1_ aTc) would allow fine tuning of the circuit output to a greater degree than with the first input alone. We created a simple *a priori* model of this circuit using nominal parameters (Figure 1A), which demonstrated that inclusion of sRNA can tune the autorepressor’s dynamic range without introducing substantial additional noise into the system’s output. To test the tunability of output, JW3876 cells containing pCK221 were grown in rich media supplemented with varying concentrations of aTc and L-rhamnose and assayed at late-exponential phase (5 h; Figure S14) as before. As predicted, a wide range of mean fluorescence values with small error bars were accessible using sRNA-tuning of the autorepressor (Figure 3C), compared to the large error bars observed with aTc-tuning alone (Figure S3), implying easier and more precise tunability. We developed a more complex four-state model of this circuit (S.I. 1.4) using some parameters from the literature with the remaining unknown parameters fitted to the experimental data using least-squares minimisation. Simulations using the updated parameters were then compared to the experimental results to test the model fit, with excellent quantitative agreement observed (Figure 3C) demonstrating that the hypotheses upon which the model was built were able to capture the observed experimental behaviour.

**Figure 3.**
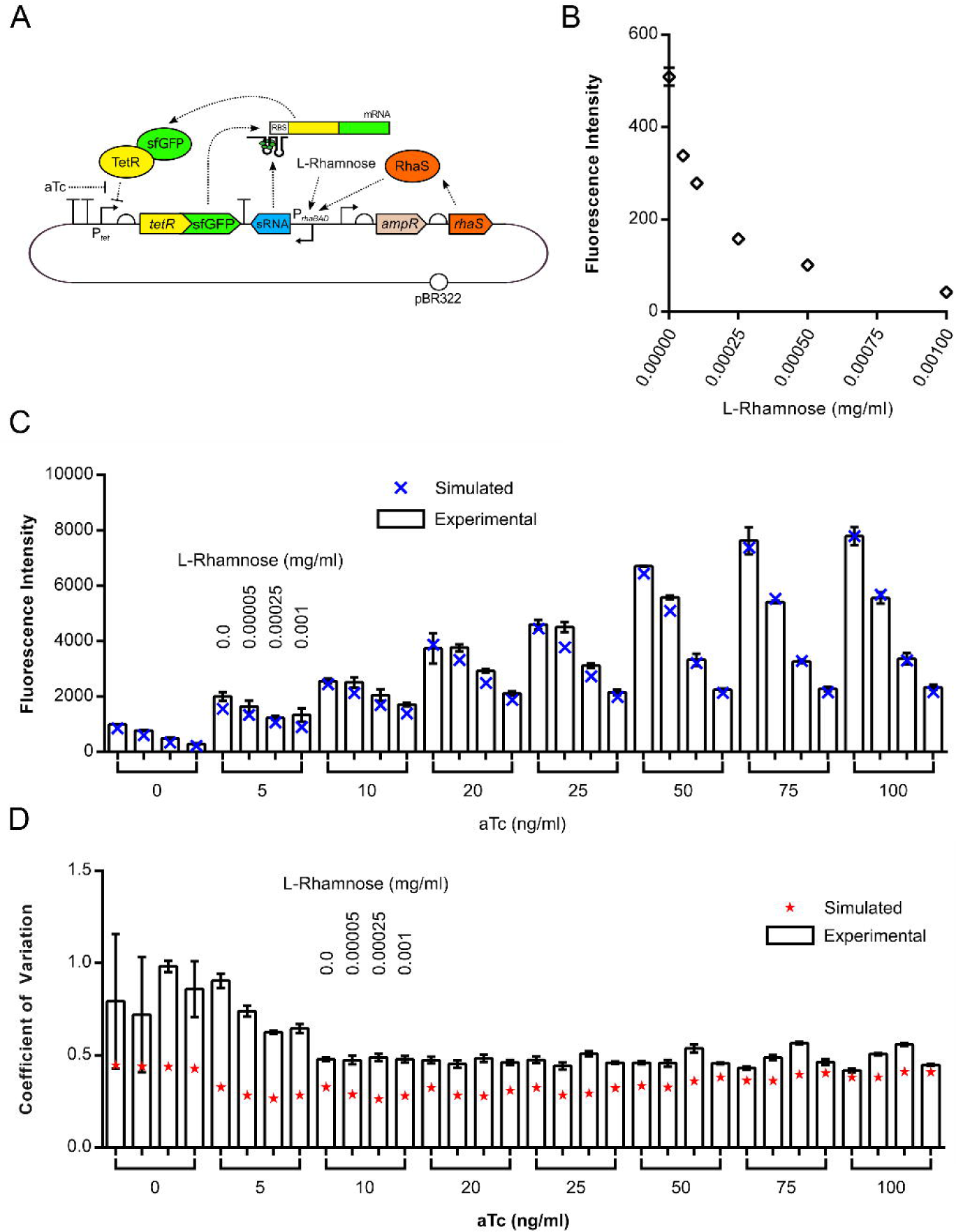
Testing the tunability of the sRNA-tuned autorepressor plasmid pCK221. A. Schematic diagram of plasmid pCK221. B. Testing the tunability of sRNA translation inhibition with increasing concentrations of L-rhamnose. *E. coli* strain JW3876 was transformed with pCK221, cultured at 37 °C in EZ rich defined medium supplemented with glycerol and increasing concentrations of L-rhamnose and GFP fluorescence measured at late exponential phase (5 h) by flow cytometry. C. Testing the ability to fine-tune the coarse aTc input-output dial with the inducible expression of the sRNA. *E. coli* strain JW3876 was transformed with pCK221, cultured at 37 °C in EZ rich defined medium supplemented with glycerol and increasing concentrations of L-rhamnose and aTc and GFP fluorescence measured at late exponential phase (5 h) by flow cytometry (white columns). Simulated steady-state output is overlayed to show model fit (blue X). D. Experimentally-obtained coefficients of variation around the TetR-sfGFP mean when output is tuned using both L-rhamnose and aTc (white columns) with predicted coefficients of variation overlayed (red stars). Fluorescence intensity represents the geometric mean of fluorescence. Error bars shown represent the standard deviation of three independent biological replicates.

Modularity is an important consideration when designing and constructing novel feedback circuits for synthetic biology. We wanted to demonstrate that this ability to tune the effective feedback strength of an autorepressor was possible using alternative inducible promoters for sRNA expression. This would allow application-specific inducible promoters, for example one that responds to an internal cellular metabolite, to be used and allow integration of the sRNA-tuned autorepressor motif into synthetic metabolic engineering projects. To this end the *rhaBAD* promoter was replaced with the *m*-toluic acid-responsive *Pm* promoter and transcriptional activator XylS from *Pseudomonas putida* (93–95) and the resulting plasmid, pCK226 (Figure S5A), assayed in rich media with a range of concentrations of *m*-toluic acid and aTc. As with the P_rhaBAD_-based plasmid, expression of the sRNA was possible with this second inducible promoter system and in combination with aTc allowed fine tuning of the sRNA-tuned autorepressor output (Figure S5C). Relevant parameters that were identified for plasmid pCK221 were carried across to modelling of pCK226, and the remaining unknown parameters identified using experimental data from pCK226. Good agreement was observed between the model and experimental data (Figure S5C), demonstrating that our model can be applied for predictive purposes even when its cellular context is changed.

The noise profile of the circuit was simulated using stochastic models to assess the contribution of intrinsic and extrinsic noise to the overall circuit noise and these were subsequently compared to experimental data (SI 1.4.5). The simulations suggest that extrinsic noise (resulting from cell-wide variations such as nutrient concentrations, ribosome numbers, inducer uptake etc.) contributes more to the overall noise in our system, than intrinsic noise (resulting from stochasticity of reactions within the system) (Figure S7). When the contribution of both types of noise are combined, the noise profile is flat with increasing concentrations of either *u*_1_ (aTc) or *u*_2_ (L-rhamnose) (Figure 3D). Coefficients of variation were extracted from the flow cytometry data for each sample and no statistically-significant difference in CoV was observed with increasing sRNA expression (*u*_2_) or *u*_1_ at concentrations above 10 ng/ml (Figure 3D), which is in good agreement with predictions. A similar noise profile was obtained with the pCK226-encoded circuit (Figure S5D & E).

### A closed-loop sRNA-based negative feedback circuit

The performance and analysis of the sRNA-tuned autorepressor demonstrates that engineered sRNAs can play a valuable role in synthetic negative feedback circuits. We next wanted to build and test a negative feedback circuit where the sRNA is in closed-loop feedback with the output of the circuit (Figure 1B). The use of an sRNA-mRNA summing junction in a translation-based feedback circuit could have many advantages over transcription-based feedback circuits such as the autorepressor, including the following. The use of a short (approximately 200 bp), double-stranded nucleic-acid fragment avoids overproduction of a protein-based controller, which can increase cellular burden and impact downstream applications of the negative feedback circuit. sRNA-based feedback is predicted to allow tighter regulation of the output as mRNAs are quickly silenced upon Hfq-mediated sRNA-mRNA binding (38). The ability to vary the concentration of an sRNA allows stoichiometric silencing of mRNAs, which allows predictable, linear tuning of the feedback strength.

A closed-loop sRNA-based negative feedback circuit was designed and simulated. It consists of a transcriptional activator that in the presence of an external input (*u*_3_) induces expression of a translation-inhibiting sRNA, which itself targets the mRNA encoding the transcriptional activator and thus reduces production of the activator protein. We investigated this circuit’s anticipated behaviour using a simple model with nominal parameter values (Figure 1B) and found that the introduction of feedback (with increasing *u*_2_) reduced the system’s output and coefficient of variation. To validate the general behaviour of this circuit experimentally, the two halves of the circuit were split across two plasmids. The first plasmid, pAH12 contains *rhaS*, the gene encoding the transcriptional activator of the *rhaBAD* promoter, fused to the gene encoding sfGFP. A plasmid backbone was chosen with a mean copy number of 17 and the strong constitutive proD promoter (96) was inserted upstream of *rhaS.* Two different synthetic sRNAs were designed as before, one targeting the start codon of the rhaS-sfGFP mRNA (Start) and the other targeting the Shine-Dalgarno of the rhaS-sfGFP mRNA (SD). Both sRNAs were analysed using IntaRNA, with the SD sRNA found to bind more favourably than the Start sRNA to the rhaS-sfGFP mRNA (−36.01 kcal/mol versus −33.53 kcal/mol respectively) and no substantial off-target binding predicted (Table S2). Each of the sRNA constructs were inserted into plasmid backbones with a plasmid copy number of 15-20 and the *P*_*rhaBAD*_ promoter inserted upstream of the sRNA construct allowing RhaS- and L-rhamnose-induced expression of each sRNA, resulting in the final plasmids pAH15 (Start) and pAH16 (SD). To test the performance of this circuit, JW3876 cells containing pAH12 and either pAH15, pAH16 or pAH23 (negative control plasmid) were grown in rich media supplemented with 0.2 mg/ml L-rhamnose in a microplate reader and optical density (measured at 600 nm) and fluorescence compared at mid-exponential phase (4 h). Both sRNA plasmids were capable of reducing the amount of RhaS-sfGFP produced (highly statistically-significant difference; *P* < 0.0001) (Figure 4A), with the SD-targeting version more effective than the Start-targeting version. The SD-targeting sRNA construct was subsequently inserted into the backbone of pAH12, resulting in plasmid pCK222 (Figure S12).

**Figure 4.**
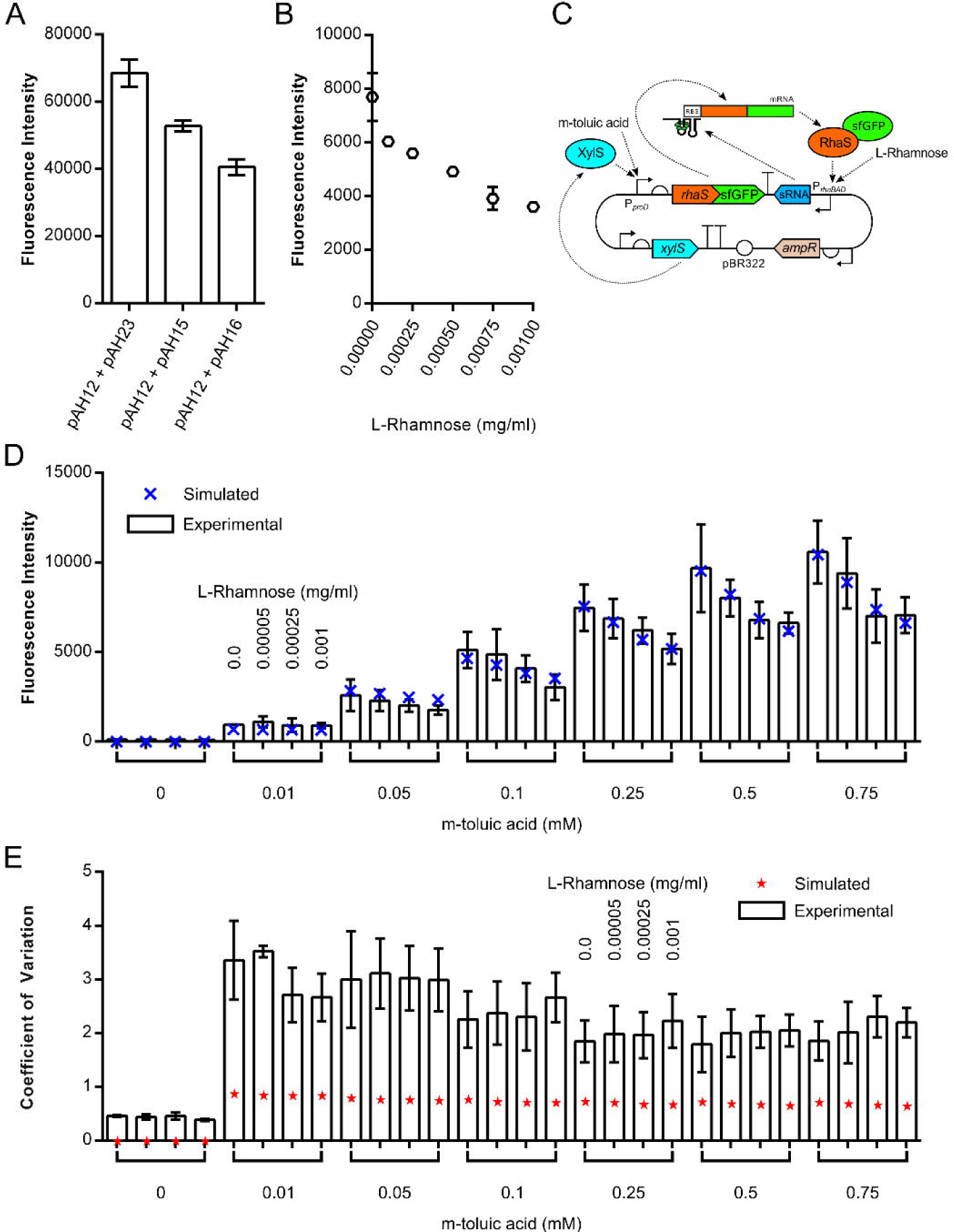
Characterisation of a closed-loop negative sRNA feedback circuit. A. Plasmids allowing the inducible expression of sRNAs targeting the Shine Dalgarno (pAH15) or start codon (pAH16) of the rhaS-sfGFP mRNA were designed and constructed and the ability of each sRNA to inhibit translation assessed. *E. coli* strain JW3876 was transformed with pAH12 (constitutive expression of *rhaS-sfGFP)* and one of pAH15, pAH16 or pAH23 (negative control, no sRNA), cultured at 37 °C in EZ rich defined medium supplemented with glycerol and 0.2 mg/ml L-rhamnose, and GFP fluorescence measured at late exponential phase (5 h) by flow cytometry. B. The tunability of sRNA translation inhibition with increasing concentrations of L-rhamnose was tested. *E. coli* strain JW3876 was transformed with pCK222 (constitutive expression of *rhaS-sfGFP* and the P_*rhaBAD*_-SD sRNA construct), cultured at 37 °C in EZ rich defined medium supplemented with glycerol and increasing concentrations of L-rhamnose, and GFP fluorescence measured at late exponential phase (5 h) by flow cytometry. C. Schematic diagram of plasmid pCK227 where the proD promoter in front of *rhaS-sfGFP* on pCK222 (Figure S12) is replaced by *xylS* and the *Pm* promoter from *Pseudomonas putida.* D. Testing the ability to set the output level with one external input and the feedback strength with another external input. *E. coli* strain JW3876 was transformed with pCK227, cultured at 37 °C in EZ rich defined medium supplemented with glycerol and increasing concentrations of L-rhamnose and *m*-toluic acid, and GFP fluorescence measured at late exponential phase (5 h) by flow cytometry (white columns). Simulated steady-state output is overlayed to test model fit (blue X). E. Experimentally-obtained coefficients of variation around the TetR-sfGFP mean when output is tuned using both L-rhamnose and *m*-toluic acid (white columns) with predicted coefficients of variation overlayed (red stars). Fluorescence intensity represents the geometric mean of fluorescence. Error bars shown represent the standard deviation of three independent biological replicates.

Next the response of this circuit to increasing concentrations of the inducer L-rhamnose was tested. Simulations predicted that in contrast to the sRNA-tuned autorepressor circuit, the response of the closed-loop sRNA feedback circuit saturates while the output is still high (Figure 1B). This results from a limiting of sRNA production, preventing complete silencing of the rhaS mRNA. To test this, JW3876 cells containing the plasmid encoding the entire circuit, pCK222, were grown in rich media with increasing concentrations of L-rhamnose and fluorescence assayed at late exponential phase (5 h) via flow cytometry. As L-rhamnose concentrations increase from 0 to 0.0008 mg/ml, an almost linear reduction in output level is observed (Figure 4B). As concentrations approach 0.001 mg/ml, saturation of the output response is observed and at the higher concentration of L-rhamnose, 0.01 mg/ml, no further decrease in output is observed (data not shown).

As the sRNA is in closed-loop feedback with the rhaS mRNA, varying the steady state of the output must be realised through changing the strength and rate of *rhaS-sfGFP* transcription. Replacing the constitutive promoter with an inducible promoter would allow the setpoint of the output to be dynamically controlled through the use of an external input, and the variability in this output controlled via the sRNA feedback loop. To this end the constitutive proD promoter was replaced with *m*-toluic acid-responsive *Pm* promoter and transcriptional activator XylS from *Pseudomonas putida* (93) resulting in plasmid pCK227 (Figure 4C). As before, JW3876 cells containing pCK227 were grown in rich media with varying concentrations of both *m*-toluic acid (inducer *u*_3_) and L-rhamnose (inducer *u*_2_) and steady state fluorescence measured at late exponential phase (5 h; Figure S14). In the absence of L-rhamnose and thus feedback, increasing concentrations of *m*-toluic acid resulted in higher output levels approaching saturation above 0.75 mM *m*-toluic acid (Figure 4D). When feedback is added via increasing concentrations of L-rhamnose, a sharp reduction in steady-state output is observed between 0 and 0.00025 mg/ml L-rhamnose followed by a gradual levelling off of output between 0.00025 and 0.001 mg/ml L-rhamnose (Figure 4D). This behaviour is consistent with the feedback-loop equilibrium being reached at the highest concentration of L-rhamnose and is accurately captured by our model, with an excellent fit to the experimental data when unknown parameters were identified (Figure 4D). The use of one external input to set the output level and another external input to tune the feedback strength allows access to a wide range of output levels, while ensuring mean output is maintained at this set level.

The noise profile of the circuit was simulated using stochastic models to assess the contribution of intrinsic and extrinsic noise to the overall circuit noise and these was compared to experimental data. As with the sRNA-tuned autorepressor results, the simulations suggest that extrinsic noise contributes more to the overall noise of a system, than intrinsic noise (Figure S11). When the contribution of both types of noise are combined, the noise profile is flat with increasing concentrations of *u*_2_ (L-rhamnose) and gradually reduces noise with increasing concentrations of *u*_3_ (*m*-toluic acid) (Figure 4E). Coefficients of variation were extracted from the flow cytometry data for each sample. No statistically-significant difference was observed with increasing sRNA expression (*u*_2_) or increasing *rhaS-sfGFP* expression (*u*_3_) (Figure 4F), both of which are in good agreement with predictions. Overall the level of noise in this circuit is higher than the sRNA-tuned autorepressor.

## DISCUSSION

In this paper we report the first use of translation-inhibiting sRNAs in synthetic negative feedback circuits. The first circuit introduces an sRNA-mRNA summing junction into the feedback loop of an autorepressor. We have shown that the level of sRNA expression can be controlled using inducible promoters and the use of this sRNA dial linearises the coarse sigmoidal input-output response of the autorepressor and its associated input, allowing more precise tuning of the circuit output. The performance of the circuit was predicted using a deterministic model and found to accurately capture the mean output of the circuit in response to both inputs. The second circuit uses translational regulation via an sRNA in the central feedback loop of the circuit. Inducible promoters are used to control the transcription of both the output-encoding gene and the sRNA, allowing two external inputs to manipulate the system. One input is primarily used to set the steady state of the circuit output and the other input used to establish feedback through sRNA expression. The use of the sRNA in closed-loop feedback ensures the desired mean output is restricted to a small range and this behaviour is in excellent agreement with simulations based on deterministic model of the circuit. In the course of revising this manuscript, the modelling of a similar sRNA-based circuit has been described, with many of the desirable attributes that have been shown experimentally in this paper simulated (97).

It has been suggested that the use of sRNAs in regulation is associated with increased noise when compared to the use of transcription factors (59). Modelling and experimental data demonstrated that the increasing sRNA expression in either the sRNA-tuned autorepressor or the closed-loop sRNA negative feedback circuit did not correspond to an increase in noise. In the case of the closed-loop sRNA feedback circuit however, an increase in noise is observed immediately upon induction of the *rhaS-sfGFP* gene, and the overall CoV levels are higher than the CoV levels of the sRNA-tuned autorepressor. This is likely due to the use of *m*-toluic acid inducer (*u*_3_), perhaps due to inconsistent uptake by all cells, which could be caused by dissociation of the organic acid at the slightly basic pH of the EZ media (approximately pH 7.4) as suggested previously (95). This hypothesis is supported by CoV data from the sRNA-tuned autorepressor circuit encoded by pCK226, where the XylS-Pm system is used to express the sRNA, where the coefficients of variation are much higher (Figure S5E) than the with the RhaS-P_*rhaBAD*_ system, pCK221 (Figure 3D).

In addition to the above, the mathematical models predicted a number of features that were not explicitly observed, but to a large extent led to interest in the circuits prior to implementation. The response time of a circuit, which we define as the time taken to reach 90% of the steady-state output, is a measure of how quickly steady-state output equilibrium is reached. Using the full four-state models and experimentally-derived parameters, step responses of each circuit were simulated when specific concentrations of each inducer were applied (Figure 5). These models assume cells are maintained in the exponential growth phase both before and after addition of inducer. Overall the time scales for both circuits to reach 90 % of steady-state output are short (1 - 2 h), though additional factors that would introduce a delay such as inducer-uptake by cells are not modelled. As mentioned, tuning the output of the P_*tet*_-TetR autorepressor through the use of aTc is difficult due to the sigmoidal shape of the input-output response curve. Step-response simulations of the autorepressor with different concentrations of aTc highlight another limitation of to this circuit architecture. As increasing concentrations of aTc are supplied, the steady-state output indeed increases, but the response time also increases proportionally (Figure 5A). Step-response simulations of the sRNA-tuned autorepressor, where aTc is fixed and expression of sRNA is increased through increasing concentrations of L-rhamnose, show no such change in response time as the steady-state output changes (Figure 5B). This demonstrates that the use of an sRNA to tune the effective feedback strength of an autorepressor allows a decoupling of response time from steady-state output. This contrasts with previously-described feedback circuits, including a recent transcriptional negative autoregulation feedback circuit, where the use of an RNA transcriptional attenuator did not allow decoupling of output and response time (98). Step-response simulations of the closed-loop sRNA feedback circuit were also performed using the full four-state models and experimentally-derived parameters. In the absence of feedback, i.e. where no sRNA is expressed (*u*_2_ = 0), response times are slower than when the sRNA feedback has been established through the supply of L-rhamnose (*u*_2_ = 2) (Figure 5C & D), where small variations in response time were observed (Figure 5D).

**Figure 5.**
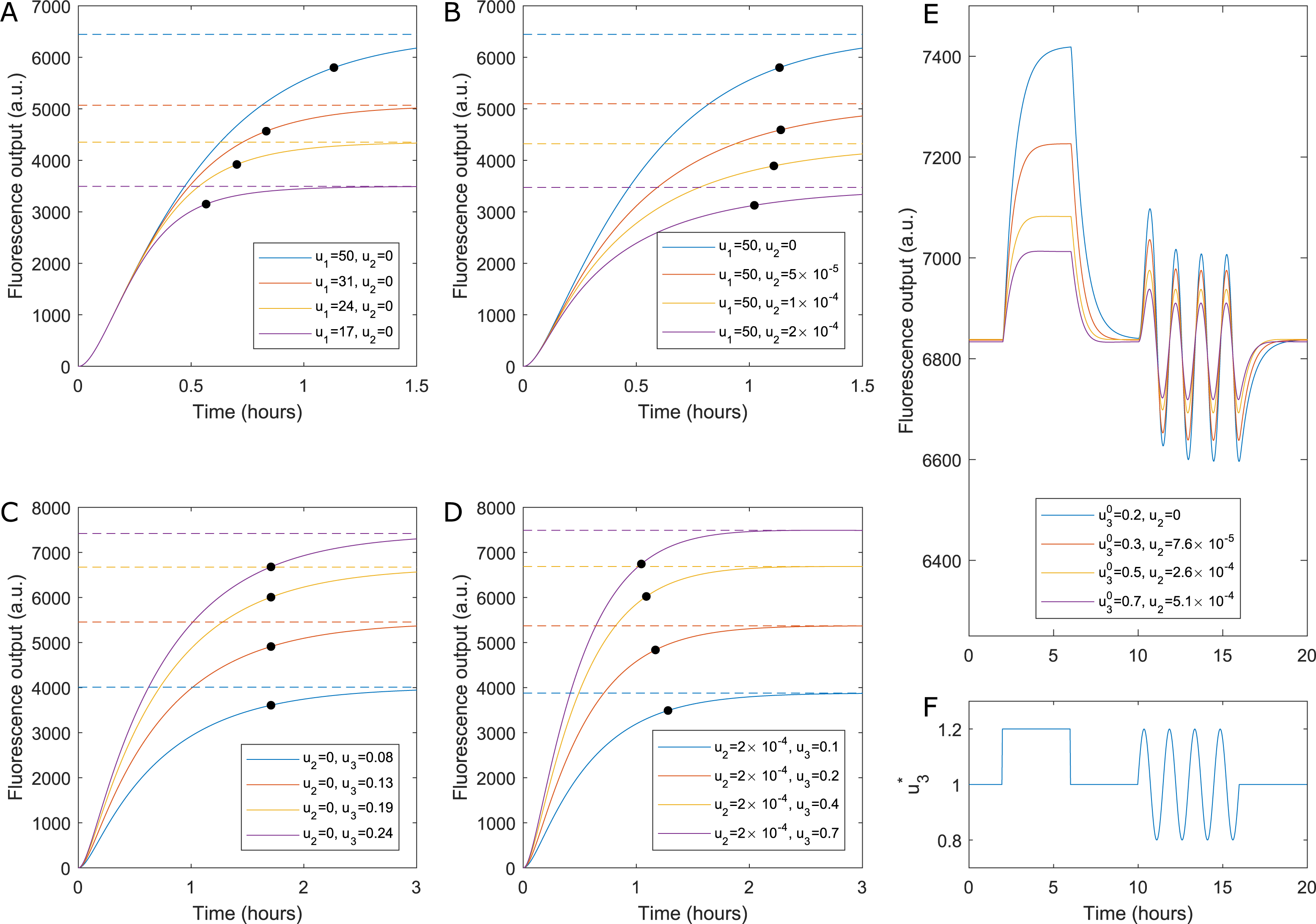
Dynamic simulations of models fit to experimental data. A. Simulation of step responses for the Autorepressor without sRNA induction. The system’s response time (black dots, defined as the point at which the output reaches 90% of its maximum value) increases as the system’s input *u*_1_ is increased. B. Simulation of the Autorepressor with sRNA induction, demonstrating that sRNA tuning allows the same absolute output levels to be achieved without substantially changing response time. C. Simulation of the closed-loop sRNA circuit without feedback (*u*_2_ = 0). D. Simulation of the closed-loop sRNA circuit with feedback (*u*_*2*_ >0), demonstrating that feedback allows the system to achieve the same output levels with a shorter response time. E. Demonstration of disturbance rejection for the closed-loop sRNA feedback circuit. A time-varying signal (sub-figure F) is applied to *u*_3_ such that 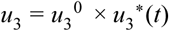. When feedback is introduced by increasing *u*_*2*_ the system’s sensitivity to an equivalent variation in *u*_*3*_ is reduced. By tuning both inputs it is therefore possible to achieve a given output level while simultaneously reducing the effect of disturbances at one input.

Disturbance rejection is another desirable characteristic of negative feedback circuits, reducing the influence that changes in input levels or the circuit environment have on their steady-state output level. For both sRNA-based feedback circuits, the models with experimentally-derived parameters were used to assess the dynamic behaviour of the circuits by numerically integrating the four differential equations governing the biochemical reactions, and applying a dynamic input profile to each inducer (SI 1.4.4 and 1.5.4). For the sRNA-tuned autorepressor, the dynamic simulations show that within a certain range of inducer concentrations, the introduction of the sRNA to the autorepressor circuit reduces the sensitivity of the circuit to fluctuations in either inducer, whilst maintaining the same mean output (Figure S6). For the closed-loop sRNA feedback the dynamic simulations show that when feedback-loop equilibrium has been reached, the output of this circuit is sensitive to changes in the first input (*u*_3_) but insensitive to changes in the second input (*u*_2_) (Figure S9). Additionally, when feedback is introduced by increasing the concentration of *u*_2_, disturbances resulting from changes in *u*_3_ concentration are dampened by the action of the feedback mechanism (Figure 5E). It is likely that other disturbances such as temperature and nutrient changes or depletion of shared cellular machinery would be rejected well by this closed-loop sRNA feedback circuit. Indeed in the course of revising this manuscript, two negative-feedback circuits using sRNAs to counter cellular burden and ribosome depletion from heterologous pathway expression have been described ((99) and doi: https://doi.org/10.1101/336271).

While the designs outlined in this paper are very simple and use well-characterised, commonly-employed components, their interconnection in a feedback fashion demonstrates the advantages of different architectures, as predicted by mathematical modelling. Although the output of both feedback circuits was a transcription factor fused to a fluorescent reporter protein, this could easily be modified to include almost any other protein of interest. Modified versions of these circuits could potentially be used to regulate and optimise the commercial production of protein therapeutics and industrially-relevant enzymes or in metabolic engineering, to ensure the concentration of the rate-limiting enzyme in a pathway is maintained. The use of negative feedback circuits has been proposed as a method to ensure the reliable interconnection of two genetic circuit modules, where the output of one circuit forms the input of another (doi: https://doi.org/10.1101/336271). The avoidance of a protein-based controller in negative feedback circuits such as these should be beneficial in these applications, through the expected reduction in cellular burden associated with using an sRNA over another resource-sapping protein. Indeed although not the focus of this work, no difference in growth was observed between each circuit in the presence or absence of sRNA (Figure S15A, B & C), implying that the sRNAs used in these feedback circuits impart a low burden upon their hosts and supporting this hypothesis. As sRNAs are simple nucleotide oligomers, with careful design they can be constructed to target almost any mRNA sequence required. Re-engineering transcription factors to bind to different DNA sequences is possible but less straightforward than redesigning nucleic acid sequences. Additionally one sRNA sequence can be designed to target multiple mRNAs with different efficiencies, opening up the possibility of synthetic genetic feedback circuits of greater complexity than has been possible to date. Moving forward it is likely that synthetic biological circuits that utilise small RNAs for control purposes, as has been demonstrated in this paper, will provide a valuable component for the engineering of biological systems to tackle a range of scientific and industrial challenges.

## ACKNOWLEDGEMENTS

The authors thank Prof. Wei Huang for hosting CK, Dr. George Wadhams and Prof. Judith Armitage for their support and advice, Dr. Tom Ellis for access to the Attune NXT flow cytometer and William Shaw and assistance with the flow cytometer.

